# Ablation of *Ezh2* in neural crest cells leads to Hirschsprung's disease-like phenotype in mice

**DOI:** 10.1101/265868

**Authors:** Hana Kim, Ingeborg M. Langohr, Mohammad Faisal, Margaret McNulty, Caitlin Thorn, Joomyeong Kim

## Abstract

In the current study, we examined the role of *Ezh2* as an epigenetic modifier for the enteric neural crest cell development through H3K27me3. *Ezh2* conditional null mice were viable up to birth, but died within the first hour of life. In addition to craniofacial defects, *Ezh2* conditional null mice displayed reduced number of ganglion cells in the enteric nervous system. RT-PCR and ChIP assays indicated aberrant up-regulation of *Zic1*, *Pax3*, and *Sox10* and loss of H3K27me3 marks in the promoter regions of these genes in the myenteric plexus. Overall, these results suggest that *Ezh2* is an important epigenetic modifier for the enteric neural crest cell development through repression of *Zic1*, *Pax3*, and *Sox10*.

## 1. Introduction

The neural crest cells (NCCs) are a multipotent cell population which have evolved among land vertebrates to develop a head, jaw, and sensory nervous system to adapt to predatory lifestyles (Gans and Northcutt, 1983). The NCCs arise between the neural epithelium and epidermis after neurulation and migrate throughout the body to generate many different cell types. Depending on their rostro-caudal axis origin and cellular differentiation capabilities, NCCs can be divided into cranial, sacral, cardiac, trunk and vagal NCCs (Bhatt et al., 2013). A number of different genes have been characterized to be important for the development of NCCs. Bone morphogenetic proteins (Bmps) and Wnt proteins induce NC formation. *Pax3* and *Zic1* specify the neural plate border (Betancur et al., 2010, Millet and Monsoro-Burq, 2012), and *Sox10* sustains NCC multipotency (Kim et al., 2014). *Hox* genes, which are critical for body segmentation and axial patterning (Maconochie et al., 1996; Holland and Garcia-Fernández, 1996), also affect the development of NCCs (Schwarz et al, 2014). Mutations in these genes can lead to neurocristopathies, which encompass a wide range of congenital diseases of neural crest cell origin. In some examples of neurocristopathies, such as Hirshsprung’s disease and Waardenburg syndrome, the enteric nervous system (ENS) is improperly developed, leading to intestinal aganglionosis in the distal portions of the colon (Moore and Johnson, 2005). Most of the ENS develops from vagal NCCs, and some contribution of sacral NCC have been seen in mice and chicks (Serbedzija et al., 1991; Burns et al., 2000; Burns and Le Douarin, 2001). Although a number of genes have been characterized to be associated with the neurocristopathies, the penetrance for the mutant allele of these genes varies suggesting a multigenic and potentially epigenetic involvement in manifestation of these disorders.

The polycomb group genes (PcGs) are epigenetic modifiers that play a significant role in regulating the expression of *Hox* genes and many other developmental transcription factors throughout embryonic development (Beuchle et al., 2001; Soshnikova and Duboule, 2009). The two major groups of PcGs include Polycomb repression complex 1 (PRC1) and Polycomb repression complex 2 (PRC2). These protein complexes are evolutionarily well conserved from flying insects to all mammals, including humans. These two complexes modify histones as an epigenetic signal: the ubiquitination on lysine 119 of histone 2B (H2BK119) by PRC1 and the methylation on lysine 27 of histone 3 (H3K27) by PRC2. In the case of PRC2, Enhancer of Zeste homolog 2 (*Ezh2*) is the histone methyltransferase, adding di- and tri-methyl groups to H3K27. The tri-methylation of H3K27 (H3K27me3) is usually associated with gene repression (Cao et al., 2002; Boyer et al., 2006; Barski et al., 2007). Complete deletion of *Ezh2* leads to early embryonic lethality (O’Carroll et al., 2001), and tissue specific deletions of *Ezh2* display loss of cellular differentiation (Zhang et al., 2015, Ezhkova et al., 2009). In particular, deletion of *Ezh2* in NCCs hinders the development of craniofacial bone structure in association with the de-repression of *Hox* genes in the first brachial arch (BA1) cells. *In vitro* culture of these BA1 cells demonstrated that differentiation factors such as GFAP and NF160 were properly expressed in *Ezh2* conditional null cells (Schwarz et al., 2014). However, these mutant NCCs have not been examined in later stages of embryonic development *in vivo*.

We performed a series of conditional knockout experiments targeting *Ezh2* to examine the effect of PRC2 in neural crest cell development in later stages of embryogenesis focusing on the ENS development. In addition to aberrant facial structures, conditional ablation of *Ezh2* in NCCs led to improper development of ganglion cells in the ENS and derepression of *Pax3*, *Zic1*, and *Sox10*.

## 2. Material and Methods

### 2.1 Animals

The *Ezh2* conditional knockout mice were derived as previously described (Su et al., 2003). Presence of a loxP site insertion was confirmed through polymerase chain reaction (PCR) amplification with the following primers: LoxF- CCCATTGAGAGTGCTGACTCA; LoxR- ACCTCGCTATGTGTAACCAGT, and F3.1- TCTTAGCACTTGCTTGTTCCCATTG. Expected fragment sizes were 100 bp without the loxP insertion and 120 bp with the loxP insertion (**Fig. S1**). Template DNA was acquired through mouse ear clips that had been digested in tail lysis buffer (0.1 M Tris-Cl, 5 mM EDTA, 0.2% SDS, 0.2 M NaCl, pH 8.0, 20 µg/ml Proteinase K). E15.5 and E18.5 embryos were harvested through timed mating between *Ezh2^flox/flox^; Wnt1-CreT/+* double mutant backcrossed with *Ezh2^flox/flox^*. The survival curve was generated by combining four litters of cesarean sections (n= 40). Pups delivered through cesarean sections were massaged on the abdomen, and fluid was gently removed from the nasal and oral cavity using Kimwipes. The p value was derived using the Fisher exact statistical test. All mouse work was approved by the IACUC committee at Louisiana State University.

### 2.2 Histology

Embryos and neonatal mice were fixed for 24-28 hours in 10% formalin and subsequently moved into 70% ethanol for storage. Tissue were trimmed and routinely embedded in paraffin, sectioned, and stained with hematoxylin and eosin (H&E).

### 2.3 Toluidine blue staining

Toluidine blue staining was performed as previously described (Spazierer et al., 2006). In brief, unfixed neonates were incubated in 100% methanol for 5 minutes and then stained in 0.1% toluidine blue dye for 20 minutes.

### 2.4 AChE staining

Acetylcholinesterase (AChE) staining was performed according to a previously described protocol (Enomoto et al., 1998). In brief, the dissected stomach and intestines were fixed in 4% paraformaldehyde for 2 hours at 4°C and moved to saturated sodium sulfate solution overnight at 4°C. Then, the tissues were incubated in the staining buffer (0.2 mM ethopropazine HCl, 4 mM acetylthiocholine iodide, 10 mM glycine, 2 mM cupric sulfate, and 65 mM sodium acetate pH 5.5) for 4-5 hours at room temperature. Lastly, acetylcholinesterase staining was completed by incubating the stomach and intestines in 1.25% sodium sulfide pH 6 for 1.5 minutes.

### 2.5 RT-PCR

The whole gut and the myenteric plexus (according to Grundmann et al., 2016) was isolated and snap frozen in liquid nitrogen. After adding Trizol (Invitrogen) into frozen tissues, the tissues were further minced into smaller fragments with sterile scissors in Trizol. Tissue fractions in Trizol were left on ice for 30 minutes and total RNA was isolated as described by manufacture’s protocol (Invitrogen). Isolated RNA was resuspended in nuclease-free water and cDNA was synthesized with MMLV reverse transcriptase (Invitrogen) according to manufacture’s protocol.

### 2.6 Chromatin immunoprecipitation assay

The P0 neonatal intestine was dissected and the myenteric plexus was isolated with Liberase TH (Roche) (Grundmann et al., 2016). Tissues were homogenized in cold 1X PBS and fixed in 0.1% PFA v/v at 37°C for 20 minutes. After three consecutive washes with cold 1X PBS, the crosslinked tissues were placed in the lysis buffer (1% SDS, 10mM EDTA, 50mM Tris-Cl pH8.1) for sonication. Chromatin fraction was pre-cleared with protein A/G agarose beads (Cat. No. sc-2003, Santa Cruz) and pre-immune serum for 2 hours. After spinning down agarose beads, the remaining supernatant was used for immunoprecipitation with H3K27me3 antibody (Cat. No. 07-449, Upstate Biotech.). The myenteric plexus isolated from two P0 neonates were combined per ChIP assay, and each ChIP assay was repeated at least three times (n=8 wild-type, n=6 Ezh2 null).

### 2.7 Whole mount RNA in situ hybridization

The E15 embryo guts were dissected and fixed in 4% PFA overnight in 4°C. The whole mount RNA *in situ* hybridization was performed as described previously (Zakin and De Robertis, 2004) with anti-Pax3 probe hybridization at 68°C. The images were captured with Olympus DP70 camera attached to the Olympus SZX7 stereo microscope.

### 2.8 Immunohistochemical procedures

The longitudinal muscle attached to the myenteric plexus (LMMP) was isolated from embryonic day 16 (E16) and P0 guts and fixed for 30 minutes in room temperature in 4% paraformaldehyde. After twenty-minute treatment in 0.05% Triton-X in 1X PBS, the LMMPs were blocked for endogenous peroxidase activity by incubation in 3% hydrogen peroxide for one hour in room temperature. The LMMPs were pre-blocked in blocking solution (1% BSA, 0.3% Triton-X in 1X PBS) for 2 hours at room temperature, and incubated in blocking solution with primary antibodies with gentle rocking at room temperature overnight. The dilutions for the primary antibody are as follows: Sox10 antibody (Santa Cruz, cat. sc365692, 1:100), GFAP antibody (Santa Cruz, cat. 33673, 1:50), HuD antibody (Santa Cruz, cat. sc28299, 1:1000). The bound antibodies were detected with Alexa Flour 488 (Invitrogen, cat. A11001, 1:200) and Alexa Flour tyramide (Invitrogen, cat. B40942) according to its manufacture’s protocols. The cells were imaged on Leica DM2500 microscope and the images were captured with 18.2 Color Mosaic camera from Diagnostic Instruments, Incorporated.

## 3. Data analysis

The current data presented were obtained from at least three biological replicates from three different mice litters and at least two experimental replicates. The survival was assessed through the Kaplan-Meier curve, and the associated p value was derived from the Fisher exact test. The p values evaluating the penetrance of the mutant phenotypes were also derived through the Fisher exact test. The statistical significance of the real-time qRT-PCR values for expression analysis and ChIP assays was evaluated using the two-tailed Student’s t-test. The neuronal cells were quantified by counting the number of neurons in 300µm^2^ area of at least eight images obtained from four mice of each wild type and mutant genotypes (n=4 wild type, n=4 *Ezh2* null). The glial cells were counted in GFAP positive cluster in 300µm^2^ area of at least eight images from four mice of each wild type and mutant genotypes. The statistical significance was tested through two-tailed Student’s t-test. The density of the nerve fibers from the AChE staining were imaged in a set magnification and was quantified through skeletal reconstruction of the ganglion cells on image J and the number of intersections were counted and divided by the relative distance analyzed.

## 4. Results

### 4.1 Conditional knockout of Ezh2 in neural crest cells leads to perinatal lethality and abnormal facial structures

To characterize the function of *Ezh2* during mouse NCC development, the mutant mice homozygous for the two loxP sites flanking exon 15 through 18 in 129P2/OlaHsd background were backcrossed with the Wnt1-Cre line in the 129/B6 mixed background, generating the *Ezh2* homozygous mice in NCC (*Ezh2^flox/flox^;Wnt1-CreT/+*; hereafter referred to as ‘***Ezh2* null**’ mice). Exon deletion and subsequent reduced expression of *Ezh2* were further confirmed through qRTPCR (**Fig. S1**). Analysis of early and late embryos from this cross showed the expected Mendelian ratios, while an increased number of dead *Ezh2* null mutants were observed among the pups found within the first day of birth (Fig. 1A, 1B, and Fig. 1C). The deletion of *Ezh2* in NCC resulted in the lethality around the time of their birth (P0).

**Figure 1.**
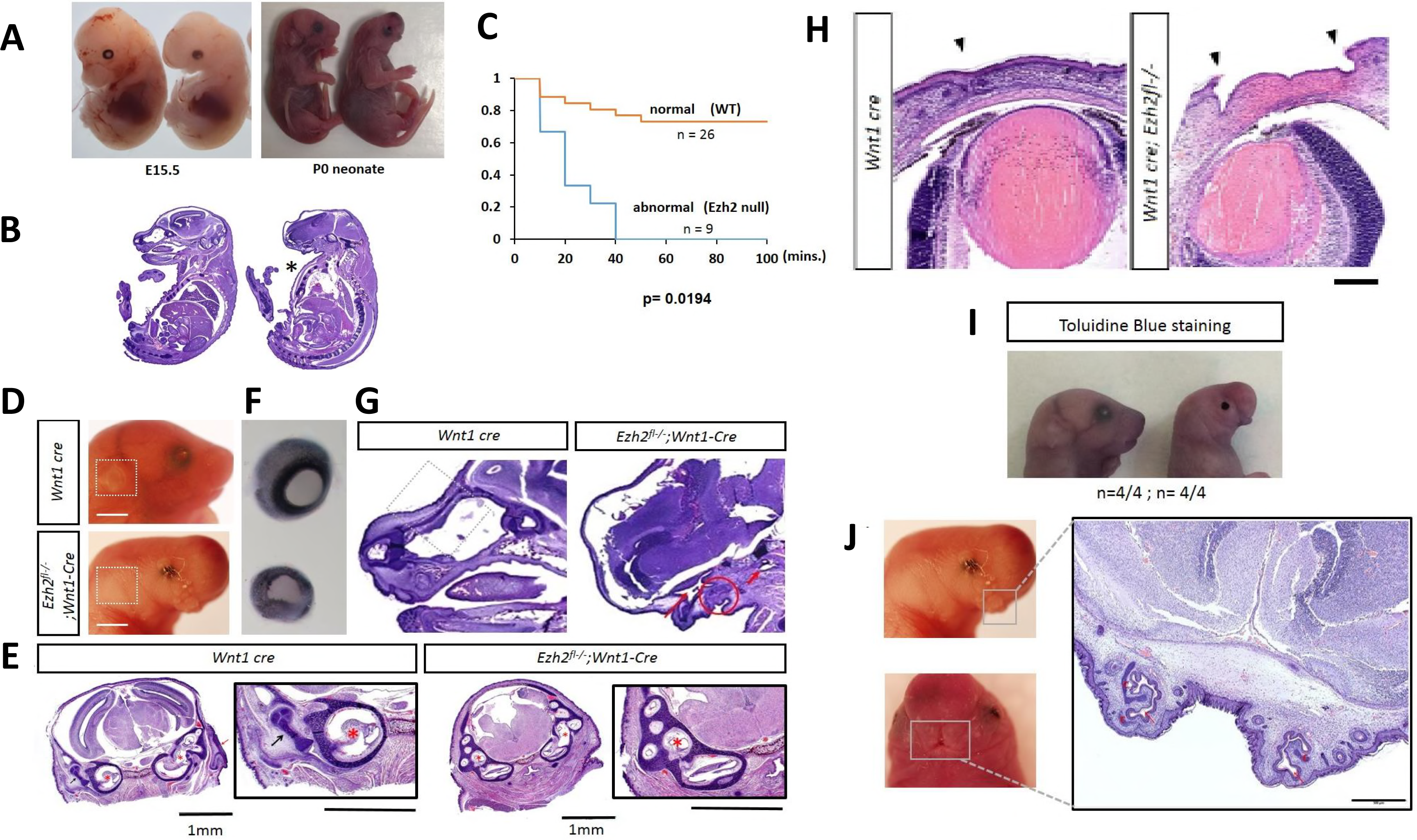
Various facial deformities and perinatal lethality in *Ezh2* null mice. **A)** *Ezh2* null mice displays abnormal craniofacial structures, which are visible at E15.5 and continues through P0. **B)** H&E staining shows absence of frontal face and ‘pigeon’s chest’ (marked in asterisk). **C)** Kaplan-Meier survival curve demonstrates perinatal lethality in *Ezh2* null mice within first 40 minutes after birth (p=0.0194, Fisher’s exact test). **D)** Absence of outer ear in *Ezh2* null mice marked in white box. **E)** Absence of the outer ear in an H&E-stained transverse section of the head of a *Ezh2* null P0 mouse, and higher magnification of an H&E-stained transverse section of the head revealing aberrant development of the middle ear in the *Ezh2* null P0 mice. *Wnt1*-Cre-Red arrow: Developing outer ear (pinna and external ear canal), Black arrow: Developing middle ear (ossicles), Red asterisks: Developing inner ear (left: cochlea; right: cochlea + semicircular canals). *Ezh2^flox/flox^;Wnt1-CreT/+***-** Red asterisks: Developing inner ear (cochlea + semicircular canals). There was no evidence of proper middle ear and outer ear development in any of the transverse sections of the head of the mutant mice. **F)** The dissected eyes of P0 pups have a 2/3 reduction in size in *Ezh2* null mutants compared to the wild-type littermates. **G)** Circle: Shows primordium of oral cavity. Long arrow: Points to the space corresponding to the nasal chamber. Short arrow: points to a space that appears to correspond to the nasopharynx (junction of nasal cavity and pharynx). **H)** Insufficient closure of the eyelid can be seen in *Ezh2* null P0 pups (marked arrowheads). H&E staining shows complete absence of cornea and conjunctiva. **I)** Toluidine blue staining of P0 pups showed staining only in the eyelid region of *Ezh2* mutants. **J)** Cleft lip (or bifid nose) visible in region of the upper lips of *Ezh2* null mice. High magnification of an H&E-stained transverse section of the head of an *Ezh2* mutant P0 mice. The arrow in red points to the nasal pits, and the surrounding cartilage plates are marked with (c) in red. The oral cavity in the center is rudimentary.

Microscopic analyses were performed to examine the facial features of the *Ezh2* null pups (P0). The results are summarized as follows. First, all of the *Ezh2* null pups displayed craniofacial defects (n=30/30), which is consistent with previous observations from the similar mutant line (*Ezh2^flox/flox^; Wnt1-Cre; Rosa26*) at E17.5 stage (Schwarz et al., 2014) (Fig. 1B and 1C). Mutant features in neonatal pups included absence of mandible (**Fig. 1G, Figs S2, S3**, and **S6)**, absence of tongue (Fig. 1G and **Fig. S6**), abnormal development of the middle ear (Fig. 1E), and absence of nasofrontal plate with herniation of the brain (frontal lobe) and meninges through the extensive cranial bone defect (meningoencephalocele) (**Fig. S2**). Second, the *Ezh2* null pups displayed absence of outer ear and microphthalmia (Figs 1D, 1F, 1B and **Fig. S2**). Consistent with the fact that the pinna develops from NCCs (Mallo, 2003), the pinnae of the outer ear were not visible in any of *Ezh2* null mice (n=30/30) (Figs 1D, 1E and **Fig. S6**). Some of the *Ezh2* null mice displayed microphthalmia: the eye was approximately 2/3 the size of that in the wild-type littermates (n=5/10) (Fig. 1F). Third, a protruding chest (or ‘pigeon’s chest’-like defect) was observed in some *Ezh2* null pups (n=4/12) (Fig. 1B asterisks). The *Ezh2* null mice also exhibit defects in closure and merging of eyelids and upper lip (Figs 1H, 1J, and **Fig. S6**). All *Ezh2* null mice (n=30/30) had incomplete development of eyelids with immature mesenchyme covered by a thin layer of epithelium remaining in their place (Figs 1D and Fig. 1H). We performed toluidine blue staining of the mice to characterize the epithelial permeability barrier. As expected, staining was detected only at the site of the non-developed eyelids of the *Ezh2* null mice (Fig. 1I, and **Fig. S5**). The epithelial barrier was intact throughout the remaining face and body of the *Ezh2* null neonatal pups, as the other areas of the body did not stain. Histologically, the *Ezh2* null neonatal pups had no cornea and conjunctiva, resulting in a malformed anterior segment of the eyes (Fig. 1H). Non-NCC lineage structures, however, such as the retina and the uvea, were similarly developed as those in the wild-type littermates (Fig. 1H). The *Ezh2* null mice also displayed incomplete fusion of the nasal placode (Fig. 1J and **Fig. S6**). Gentle massaging of the stomach in *Ezh2* null mutants after caesarian section released amniotic fluid from the mouth as it did in their wild-type littermates. This suggests that the oral cavity and the gastrointestinal tract are properly connected in the mutants. All mice, including the *Ezh2* null mutants, also had well developed lungs, indicating adequate communication of the lower respiratory tract with the nasopharynx since fetal breathing of amniotic fluid is required for proper lung development. Overall, conditional deletion of *Ezh2* in NCCs resulted in prenatal lethality and severe craniofacial defects.

### 4.2 Reduced number of ganglion cells in the enteric nervous system of Ezh2 null mice

The ganglion cells of the enteric nervous system (ENS) are also of NCC lineage. To examine proper development of ENS, series of AChE (acetylcholinesterase) histological staining were performed with the *Ezh2* null mutants and the wild-type littermates (Enomoto et al, 1998). The earliest visible difference in the morphology of these ganglion cells and nerve fibers through AChE staining was after E18.5, but was the most prominent in neonatal (P0) pups. Complete depletion of ganglion cells (aganglionosis) was visible in the distal colons of some *Ezh2* null mice (n=3 of 7 *Ezh2* null mice) (Fig. 2 – bottom panel) while milder reduction of ganglion cells (hypoganglionosis) were apparent in the other *Ezh2* null mice (n=4 of 7 Ezh2 null mice) (Fig. 2 – middle panel). In the proximal colon, approximately 90% and 60% reduction of ganglion cells in comparison to its wild type littermate were displayed from the severe and mild cases of hypoganglionosis, respectively (p<0.0001) (Fig. 2). In addition, the reduction of the enteric nervous system plexus was also observed in the small intestine and stomach of *Ezh2* null mutants compared to those in the wild-type littermates (approximately 50% reduction, p=0.0011) (**Fig. S4**). In sum, some of the *Ezh2* null mice displayed aganglionosis in the distal colon and the hypoganglionosis was extended to the small intestine and the stomach.

**Figure 2.**
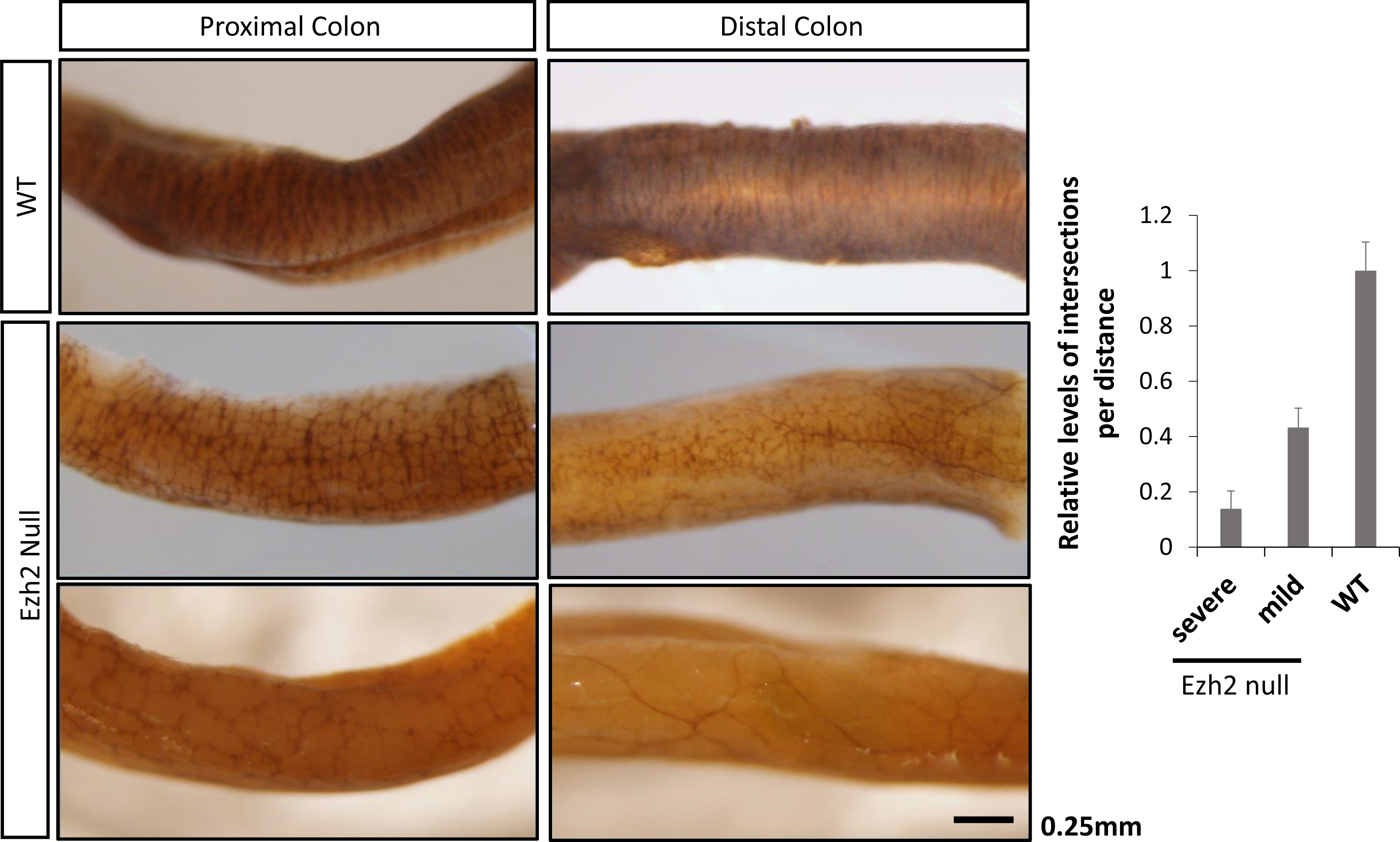
Hypoganglionosis in *Ezh2* null mice. Whole-mount AChE staining in wild type and *Ezh2* null E18.5 mice. AChE stained proximal and distal colon of wild-type littermate (top panel) and two *Ezh2* mutants (bottom panel). The two Ezh2 null mice are representative of mild (middle panel, p<0.0001) hypogangliosis and severe (bottom panel, p<0.0001) aganglionosis cases. The bar graph is the quantification of the AChE staining in the proximal colon. (scale bar = 0.25 mm).

### 4.5 Expression level changes in the whole gut of Ezh2 null mice

The effects of the *Ezh2* deletion on the enteric nervous system were further analyzed at the molecular level in the following manner. First, we examined the expression levels of NCC developmental genes using the total RNA isolated from the guts of the *Ezh2* null embryos and wild-type littermates at E18.5. Since previous studies observed over-expression of *Hox* genes in the BA1 cells of the *Ezh2* null mutants, we also tested the expression levels of *HoxA9*, *HoxA1*, *HoxA4*, *HoxA5*, *HoxB3*, and *HoxB5* (**Fig. S5**). According to the results, the majority of the tested NCC genes were down-regulated in the *Ezh2* null embryos compared to their wild-type littermates (Fig. 3). The most significant and consistent down-regulation was observed in *Phox2b* (n=4, p=0.0035) and *Sox10* (n=8, p=0.0239). In contrast, two genes, *Zic1* (n=6, p=0.003) and *Pax3* (n=6, p=0.019), were 5 to 30 fold up-regulated in the *Ezh2* null embryos compared to their wild-type littermates (Fig. 3). A slight up-regulation of *HoxA9* and *HoxA5* was seen in some of the *Ezh2* null mutants (n=2), but the levels were not as significant as *Pax3* or *Zic1* (**Fig. S5**). Gel electrophoresis of the PCR product from whole gut cDNA confirmed the presence of *Pax3* (226 bp) and *Zic1* (235 bp) expression in *Ezh2* null mutants and absence of expression in its wild-type littermates (**Fig. S7A**). To examine the aberrant spatiotemporal gene expression of *Pax3*, series of whole mount RNA *in situ* hybridization were performed on the E15 gut. All wild-type embryo guts did not show staining with the anti-Pax3 probe (n=6), while *Ezh2* null embryo showed staining throughout the entire gut (n=3) (**Fig. S7B**).

**Figure 3.**
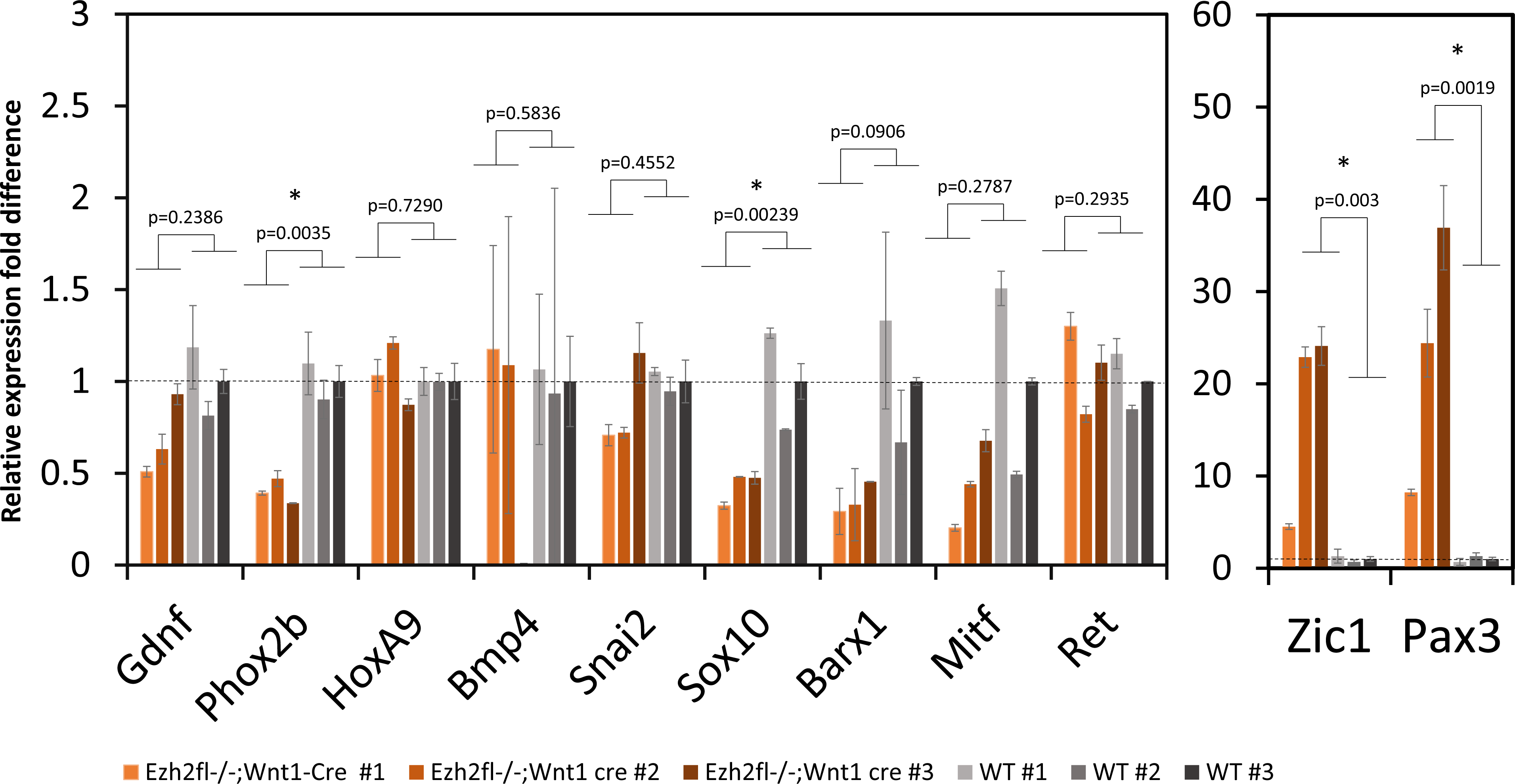
NCC developmental gene expression in *Ezh2* null mice guts. NCC developmental gene expression levels were compared between wild-type (grey) and *Ezh2* mutant (red) gut tissues with qRT-PCR. Each bar graph represents a biological replicate.

### 4.6 Expression analysis and epigenetic changes in NCC genes of Ezh2 null mice in isolated myenteric plexus

To test if the changes in the expression of these NCC developmental genes are in the enteric nervous system, the myenteric plexus was isolated from P0 neonates in accordance with previously published protocols by Grundmann et al., 2015 (Fig. 4A). The expression of *Pax3* (n=3, p=0.0010) and *Zic1* (n=3, p=0.0071) was consistently derepressed in the myenteric plexus of the *Ezh2* null mice, while no expression was observed in the wild-type littermate control. In contrast to the qRT-PCR data from the whole gut, in which the *Sox10* was seen to be down-regulated in the *Ezh2* null mice, a 20-fold up-regulation of *Sox10* (n= 4, p=0.0263) was detected in the *Ezh2* null myenteric plexus cells. Overall, *Pax3*, *Zic1* and *Sox10* are aberrantly expressed in the isolated *Ezh2* null myenteric plexus cells (Fig. 4B).

**Figure 4.**
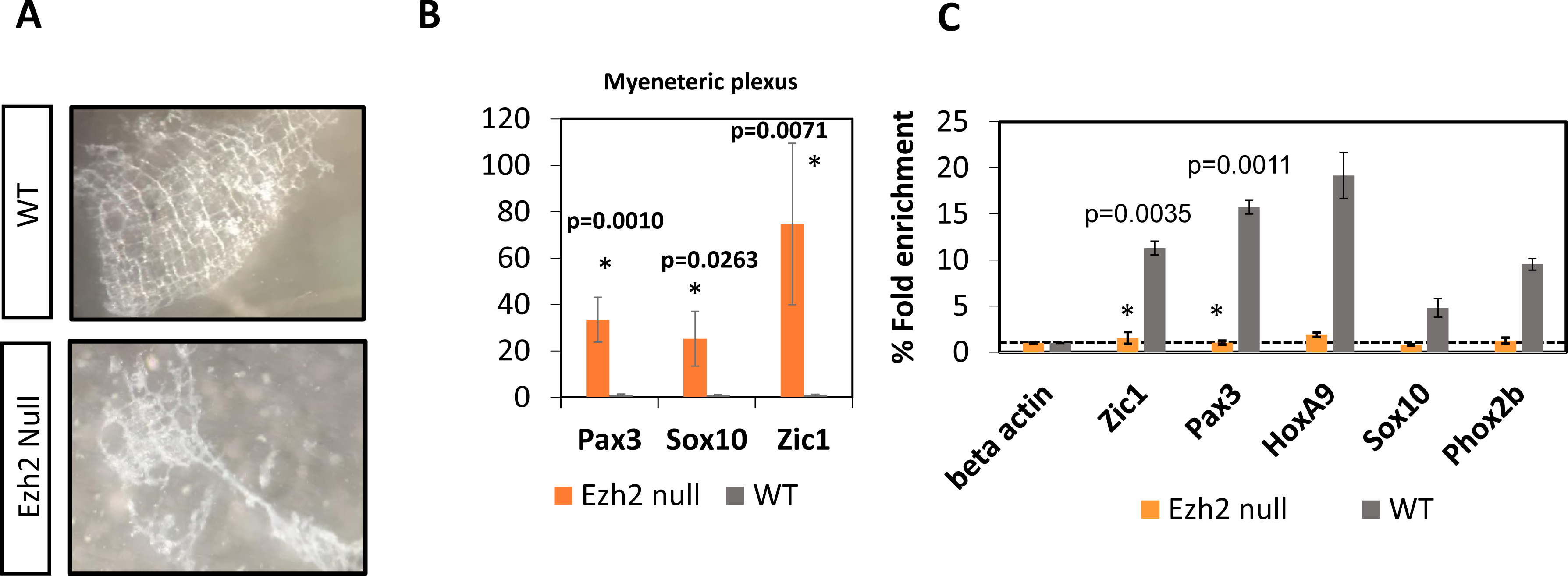
NCC gene expression analysis and H3K27me3 ChIP assays in myenteric plexus. **A)** Dissection and isolation of myenteric plexus from P0 wild-type and Ezh2 null mice for qRTPCR and ChIP assays. **B)** qRT-PCR of *Pax3*, *Zic1* and *Sox10* in myenteric plexus. **C)** H3K27me3 ChIP-qPCR in *Ezh2* mutant (in orange) and wild-type littermate (in grey) P0 gut tissues. The amount of precipitated DNA in promoter regions of NCC developmental genes and *HoxA9* are presented as a relative value (%) to that of the input DNA normalized over a negative locus (beta actin) (y-axis). Student’s t-test (two-tailed) value shows significant difference in H3K27me3 enrichment between Ezh2 null and wild-type littermates (p<0.01).

Since the H3K27me3 histone modification mark is established through *Ezh2* and this modification is associated with gene repression, ChIP assays with H3K27me3 antibodies were performed on the isolated myenteric plexus. These experiments utilized the chromatin isolated from the myenteric plexus of *Ezh2* null and wild-type littermates at E18.5 (Fig. 4A). According to the results, *Pax3* and *Zic1* showed the most significantly (p=0.0011 and p=0.0035, respectively) reduced levels of H3K27me3 enrichment in the *Ezh2* null embryos (n=6) compared to those from the wild-type littermates (n=8) (Fig. 4C). In sum, this series of analyses indicated up-regulation of *Zic1*, *Pax3* and *Sox10*, and reduced levels of H3K27me3 modifications in the promoter regions of NCC developmental genes.

### 4.7 Reduced neuronal and glial cells and aberrant accumulation of Sox10 in Ezh2 null myenteric plexus

To examine the number of neurons and glial cells in the myenteric plexus, the longitudinal muscle attached to myenteric plexus (LMMPs) was isolated from the ileum and the proximal colon and were stained with a neuronal marker Hu (Furness et al., 2004) and a glial marker GFAP. The reduced number of neuron and glial cells were visible from E16 embryos (Fig. 5), and this pattern of reduced neuronal and glial cells in the *Ezh2* null mice continued in P0 neonates (Fig. 6 and Fig. 7). On average, approximately 200 neuronal cells were counted per 300µm^2^ area in wild type P0 mice while approximately half of that amount of neurons in the same area was seen in the *Ezh2* null myenteric plexus ranging from 39 to 138 neuronal cells (n=8, p<0.001) (Fig. 7). Immunohistochemistry of GFAP also showed much less amount of GFAP positive cells in the *Ezh2* null mice compared to its littermate control in both E16 and P0 stage myenteric plexuses (n=8, p<0.0001) (Fig. 5 and Fig. 7).

**Figure 5.**
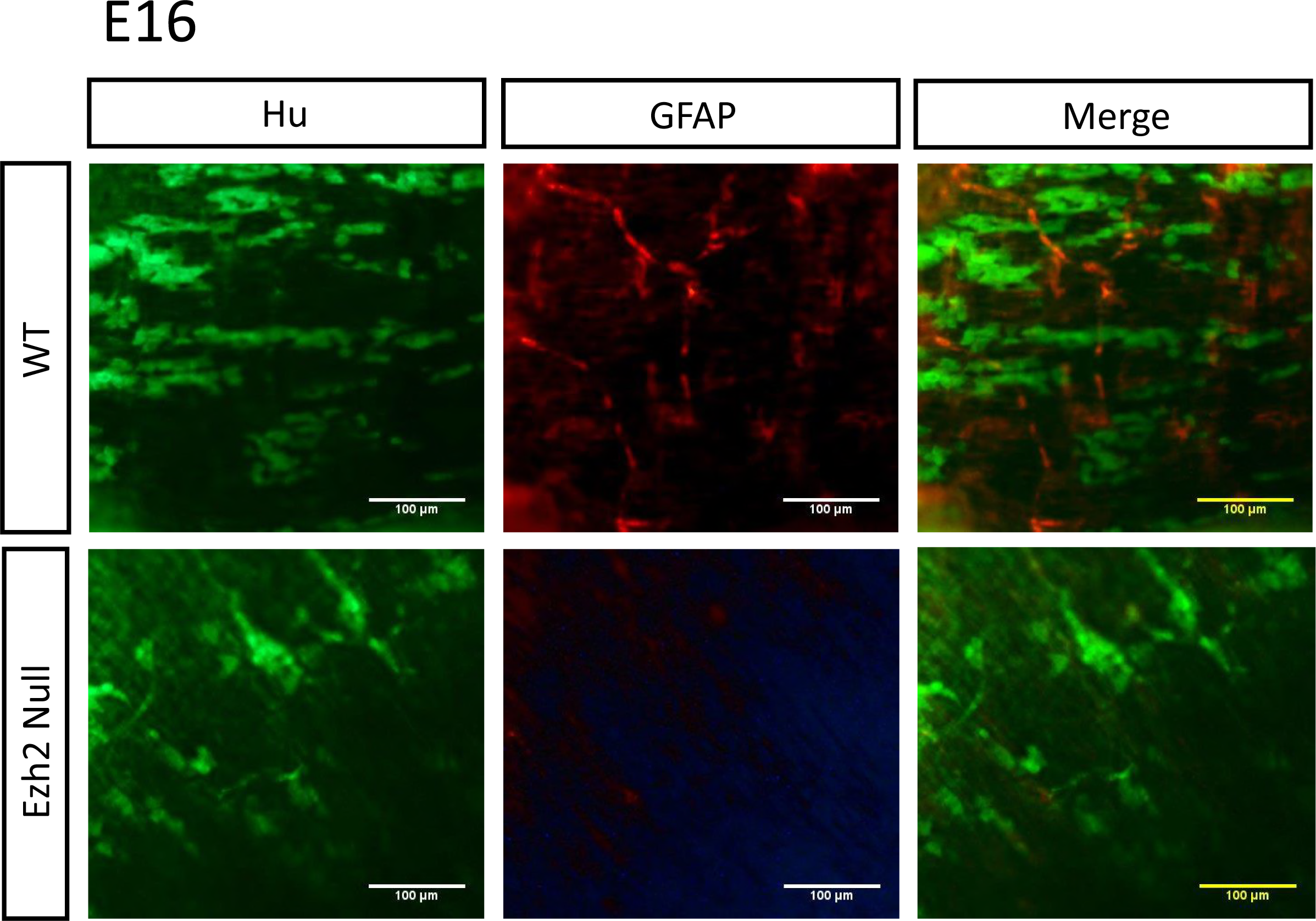
Immunohistochemistry of Hu and GFAP in E16 *Ezh2* null mice myenteric plexus. The distribution of Hu expression (green) and GFAP expression (red) in wild type (top) and *Ezh2* null (bottom) E16 myenteric plexus (scale bar = 100 µm).

As *Sox10* is an early precursor marker for NCCs (Bondurand and Sham, 2013), and it is one of the genes which displayed contradicting results in the expression patterns of the whole gut compared to the isolated myenteric plexus (Fig. 3 and Fig. 4), the expression patterns of Sox10 in the myenteric plexus of P0 stage neonates were examined with immunohistochemistry. During the enteric nervous system development, *Sox10* is expressed all migrating progenitors of ENCCs, and as the cells differentiate into neuronal lineage, *Sox10* is repressed in the neuronal cells and the expression of *Sox10* remains only in the glial cells of the developed enteric nervous system (Kim et al., 2003, Bondurand and Sham 2013). As expected, the Sox10 immunohistochemistry staining in *Ezh2* wild type control mice also did not show much overlapping expression of Sox10 and Hu (**Fig. 6 –** top panel), and almost complete overlapping expression of Sox10 and GFAP (Fig. 7– top panel). On the other hand, the *Ezh2* null mice displayed aberrant overexpression of Sox10 proteins, some of which overlapped with Hu (Fig. 6, bottom panel, white arrow) and some of which overlapped with GFAP (Fig. 7 – bottom panel, white arrow). Other clusters of Sox10 overexpressing cells did not overlap with any of these markers (Fig. 6, and Fig. 7). Overall, *Ezh2* null mice displayed reduced number of neuronal cells and glial cells in the myenteric plexus of E16 embryos and P0 stage neonates, and overexpression of Sox10 proteins were detected in the myenteric plexus P0 stage neonates of *Ezh2* null mice.

**Figure 6.**
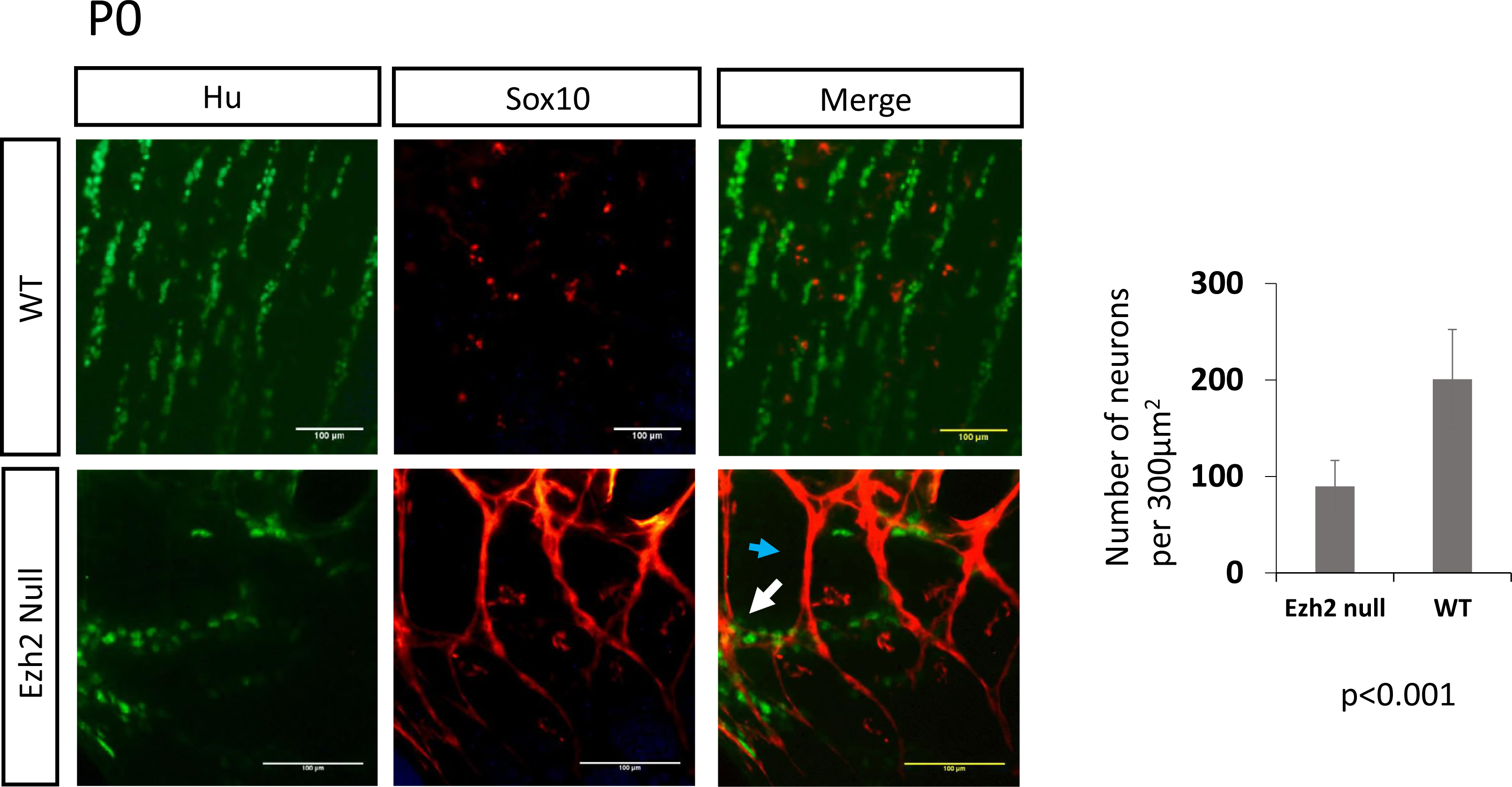
Double immunofluorescence of Sox10 and Hu in myenteric plexus of P0 neonates. Immunohistochemical analysis of Sox10 (red) in P0 neonates show non-overlapping expression of Sox10 (red) and Hu (green) in wild-type littermates (top), while *Ezh2* null mutants display aberrant overexpression of Sox10 proteins that partially overlap with Hu expression (bottom) (overlap of Sox10 and Hu marked with white arrow) (scale bar = 100 µm).

**Figure 7.**
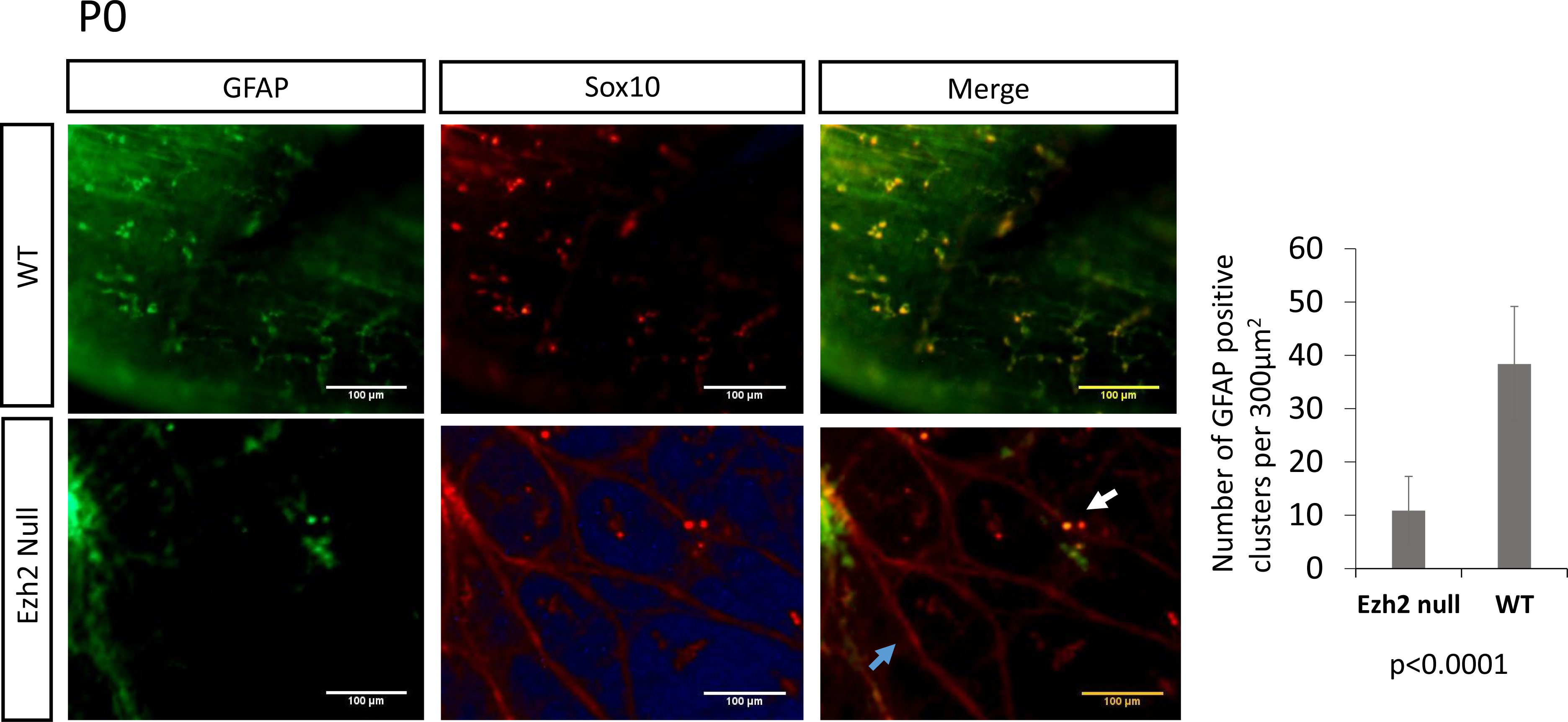
Double immunofluorescence of Sox10 and GFAP in myenteric plexus of P0 neonates. Co-expression analysis of Sox10 (red) and GFAP (green) in wild type (top panel) and *Ezh2* null (bottom panel) myenteric plexus (scale bar = 100 µm).

## 5. Discussion

In the current study, we characterized the function of *Ezh2* as an epigenetic modifier in the development of cranial NCCs and ENCCs. First, we described the mutant phenotypes exhibited by the deletion of *Ezh2* in NCCs. Then, we performed qRT-PCR, ChIP assays, and immunohistochemistry in the neonate myenteric plexus to test potential changes in the expression and epigenetic modification levels in NCC developmental genes in the *Ezh2* null mice. The results suggest a role for *Ezh2* as an epigenetic modifier for the development of ENCC in regulating the repression of *Pax3*, *Zic1*, and *Sox10* during the development of the ENCCs.

The *Ezh2* null mice were viable up to birth, but died shortly after (Fig. 1). Consistent with the previous observation, these mice displayed multiple facial deformities, which include the absence of the jaw bones, underdevelopment of the middle ear, absence of Meckel’s cartilage, absence of nasofrontal plate with herniation of the brain and meninges (Fig. 1) (Schwarz et al., 2014; Zemke et al., 2015). Some of the newly described cranial NCC mutant phenotypes include absence of outer ear, incomplete development of the eyelids, absence of the cornea and conjunctiva, microphthalmia, and bifid nose (Fig. 1). In addition to the cranial NCC defects, aberrant development of ENCCs were also often seen in *Ezh2* null mice (Fig. 2). The *Ezh2* null mice displayed hypogangliosis in later stages of embryonic development (E18.5), characterized by hypoganglionosis in the distal colon and 30%-60% reduction in the stomach and small intestine enteric plexus in *Ezh2* mutants (Fig. 2 and **Fig. S4**). Gene expression analysis in the E18.5 whole gut of Ezh2 null mice showed significant reduction of *Sox10* and *Phox2b* expression and 5 to 30 fold up-regulation of *Pax3* and *Zic1* expression compared to its wild-type littermates (Fig. 3). The spatial temporal expression of *Pax3* was confirmed through whole mount *in situ* hybridization of E14.5 gut (**Fig. S7B**) and the expression of the aberrant transcripts of *Pax3*, *Zic1* and *Sox10* were further confirmed in the isolated myenteric plexus cells (Fig. 4). In the myenteirc plexus, *Sox10*, which was down regulated in the whole gut seemed to also be surprisingly upregulated (Fig. 4). The Immunohistochemistry of *Ezh2* null myenteric plexus from E16 and P0 suggested reduced levels of both neuronal and glial cells and aberrant overexpression of Sox10 in P0 neonates (Figs 5, 6, and Fig. 7).

Two possible explanations can be suggested for the outcome of the hypoganglion mutant phenotype. First, the differentiation capacity of the ENCC precursor cells in *Ezh2* null mice may be lost due to de-repression of *Pax3* and *Zic1*. In Xenopus embryos, the over-expression of *Pax3* and *Zic1* was shown to induce the ectopic expression of neural crest differentiation downstream genes in the ventral region (Sato et al., 2004, Hong and Saint-Jeannet, 2007). During ENCC development, the neural plate border genes may need to be repressed through *Ezh2*, hence the loss of *Ezh2* in these cells might lead to the aberrant expression of *Pax3* and *Zic1*, resulting in competition for transcription factors or changes in chromatin structures of downstream gene enhancers. In the early NCC induction stage, *Pax3* and *Zic1* expression may have never been shut down due to the loss of H3K27me3 repression signals in the *Ezh2* mutants. Aberrant expression of *Pax3* in the later stages of NCC development has been reported to manifest cleft palate and abnormal osteogenesis in mice (Wu et al., 2008). *Zic1* has also been proposed to be repressed in migrating NCCs (Sun Rhodes and Merzdorf, 2006). However, the derepression of *Pax3* and *Zic1* in the *Ezh2* null mice may be limited to the ENCCs, since significant upregulation of these genes were not detected in the BA1 cells of *Ezh2* null mice in previous studies (Schwartz et al., 2014). Second, in addition to *Pax3* and *Zic1*, *Sox10* was also significantly up-regulated in the myenteric plexus cells. The conflicting results of *Sox10* expression in the whole gut versus the myenteric plexus may be due to the overall reduced amount of NC specifier genes in the gut, but increased amount of undifferentiated cells in the myenteric plexus. Notably, aberrant expression of *Sox10* has been linked to hypogangliosis in human patient studies and in mouse models (Sham et al., 2001; Southard-Smith et al., 1998; Herbarth et al., 1998; Nagashimada et al., 2012). The immunohistochemical staining of Sox10 in *Ezh2* null mice seem to favor this scenario in that the clusters of Sox10 overexpressing cells overlap with neither neuronal nor glial cells (Fig. 6 and Fig. 7) and seem to have adopted its own unique cell fate.

It is also possible that *Hox* genes are epigenetically altered in the ENS of the *Ezh2* mutants, since *Hox* genes has been linked to megacolon phenotypes (Tennyson et al., 1993), and combined mutations in *Ret* and *Hox* genes have been previously linked to increased penetrance of Hirschsprung’s disease (Garcia-Barceló et al., 2007). However, in the current studies, we have seen up-regulation of *Pax3* and *Zic1*, but not *HoxA9*, which was one of the most significantly up-regulated genes in the BA1 cells of *Ezh2* null mutants (Schwarz et al., 2014). It is interesting that the upregulated genes between the cranial NCCs are not shared in the ENCCs. The transcriptional levels of other *Hox* genes, *HoxA1*, *HoxA4*, *HoxA5*, *HoxB3*, and *HoxB5*, which are involved in the enteric nervous system development (Aubin et al., 1999, Chen et al., 2005, Tennyson et al., 1993, Fu et al., 2003, Kam et al., 2014), were also tested. However, their gene expression levels were not significantly altered in the *Ezh2* null mice (**Fig. S5**). Although ChIP assays suggest reduced H3K27me3 mark in *HoxA9* promoter region, it is puzzling to see no changes in gene expression. A number of factors have been suggested to play a role in affecting the transcriptional machinery of PRC2, including how the PRC2 subunits such as Suz12, Eed, and Ezh1 are assembled, and how these complexes interacts with other cofactors such as *Jarid2* and *Aebp2* (Aranda et al., 2015). In addition, the removal of H3K27me3 alone have been demonstrated be insufficient to activate its downstream genes (Kashyap et al., 2011). Future studies in comparing the *Ezh2* null cranial NCCs to its ENCCs and identifying the types of transcription factors that may respond to the loss of H3K27me3 and recruit gene expression machinery to its downstream genes may help to characterize the mechanism in how *Ezh2* affects NCC development.

NCCs played a significant role in vertebrate evolution, including adaptation to the environment through formation of the head, and advancement in sensory function as well as complex signaling between the gut and the brain (Gans and Northcutt, 1983; Gershon, 1997). Further understanding the conserved transcriptional regulatory mechanisms involved in the differentiation of cranial NCCs compared to vagal and sacral NCCs may help us to understand the process by which these mutipotent cell lineages have evolved. Moreover, characterizing the epigenetic modifications involved in regulating the spatiotemporal expression of these transcription factors can add to another layer of deciphering the gene regulatory network responsible for NCC development. In this study, we show the importance of *Ezh2*’s role as an epigenetic modifier in cranial and non-cranial NCC development, and we propose the involvement of the H3K27me3 epigenetic mechanism in repressing some of the neural crest cell developmental genes which can manifest a Hirschsprung’s disease-like phenotype.

## Acknowledgements

This work was supported by the National Institute of Health [J.K. R01-GM066225, R01-GM097074]. We thank the Kim lab members for their generous support and feedback in constructing the manuscript, and David Donze for sharing reagents and equipment needed in our studies.

## Author Contribution

H.K., I.M.L., and J.K. wrote the manuscript text; H.K., I.M.L., and M.F. prepared figures 1-5; M.M. and C.T. prepared supplemental figures 4-5. All authors reviewed the manuscript.

## Additional Information

**Competing financial interests:** The authors declare no competing financial interests.

